# Topological Reinforcement as a Principle of Modularity Emergence in Brain Networks

**DOI:** 10.1101/408278

**Authors:** Fabrizio Damicelli, Claus C. Hilgetag, Marc-Thorsten Hütt, Arnaud Messé

## Abstract

Modularity is a ubiquitous topological feature of structural brain networks at various scales. While a variety of potential mechanisms have been proposed, the fundamental principles by which modularity emerges in neural networks remain elusive. We tackle this question with a plasticity model of neural networks derived from a purely topological perspective. Our topological reinforcement model acts enhancing the topological overlap between nodes, iteratively connecting a randomly selected node to a non-neighbor with the highest topological overlap, while pruning another network link at random. This rule reliably evolves synthetic random networks toward a modular architecture. Such final modular structure reflects initial ‘proto-modules’, thus allowing to predict the modules of the evolved graph. Subsequently, we show that this topological selection principle might be biologically implemented as a Hebbian rule. Concretely, we explore a simple model of excitable dynamics, where the plasticity rule acts based on the functional connectivity between nodes represented by co-activations. Results produced by the activity-based model are consistent with the ones from the purely topological rule, showing a consistent final network configuration. Our findings suggest that the selective reinforcement of topological overlap may be a fundamental mechanism by which brain networks evolve toward modular structure.

## Introduction

Modularity, the presence of clusters of elements that are more densely connected witch each other than with the rest of the network, is a ubiquitous topological feature of complex networks and, in particular, structural brain networks at various scales of organization [1].

Modularity was among the first topological features of complex networks to be associated with a systematic impact on dynamical network processes. Random walks are trapped in modules [2], the synchronization of coupled oscillators over time maps out the modular organization of a graph [3] and co-activation patterns of excitable dynamics tend to reflect the graph’s modular organization [4-6]. At an abstract level, modularity in the brain is thought to be important for information processing, the balance segregation and integration as well as system evolvability in the long temporal scale, among others [1]. More concretely, the modular organization of brain networks forms the substrate of functional specialization (e.g., sensory systems [7]), contributes to the generation and maintenance of dynamical regimes (e.g., sustained activity [8] and criticality [9]), and supports the development of executive functions [10]. Thus, modularity is a key component of structural brain networks with important functional consequences.

While a number of potential mechanisms have been proposed for the creation of modules [11-13], the fundamental generative principles of the emergence of brain modules remain elusive, both algorithmically, in terms of the necessary topological changes for generating them, as well as with respect to a plausible biological implementation, that is, the realization of such topological changes through physiological mechanisms.

Generative models constitute a common approach to the study of the formation of global patterns of brain connectivity [14], where, broadly speaking, networks are allowed to grow in size and/or density according to specific rules. These models might be either based on theoretical assumptions, such as developmental time windows [15] and non-linear growth [16], constrained by experimental criteria, for instance, including geometric and topological features found in empirical connectivity data [17], or based on dynamical factors, such as the level of synchronization between nodes [18]. Given the well accepted role of synaptic plasticity in brain development and activity-dependent adaptation [19], other perspectives focus on changes driven by such local plasticity mechanisms in physiologically more realistic models. A considerable proportion of this work aims at explaining empirically observed distributions of physiological parameters at the cellular scale, such as synaptic weights [20], and only a few studies have paid attention to topological aspects, such as the proportion of local motifs [21]. Some of the mentioned modeling studies showed an emergence of modular network structure and attempted to provide an underlying mechanism based on the reinforcement of paths between highly correlated nodes [22]. Yet, the problem of a topological gradient, along which network changes should occur during the rewiring process in order to promote the emergence of modules, was not explicitly investigated.

Addressing this challenge, the present study proposes a generative principle of structural modular networks through topological reinforcement (TR). This rewiring rule, derived from a purely topological perspective, constitutes a plausible underlying mechanism leading to the formation of modules. Fundamentally, this rewiring mechanism is based on the topological overlap (TO) [23]. The origin of the TO concept stems from applications of set theory to nodes graph in network analysis, which became established as a relevant approach for quantifying the similarity of nodes in terms of their common network neighborhoods; for instance, for a review focusing on bipartite graphs see [24]. TO is closely related to the matching index [7,25], see also [26,27], an adaptation of the Jaccard index to neighborhoods of nodes in a graph. Higher-order variants of this quantity have also been discussed in the literature [28].

Prompted by the exploration of network motifs (that is, few-node subgraphs which are often statistically enriched in real networks, see [29,30]) the interplay of different topological scales in a graph has become an object of intense research. In particular, several studies have shown that global network properties, such as hierarchical organization [31] or modularity [32], can systematically affect the composition of networks in terms of local topology or network motifs, see also [33]. Intriguingly, that line of research inspires the complementary possibility: a systematic iterative selection on local network structures may conversely install, or at least enhance, certain global network properties. This is the conceptual approach we set out to explore here, where our topological reinforcement rule iteratively enhances the local topological overlap.

As a further step, we explore a plausible dynamical implementation of the topological reinforcement. We used an excitable network model, the SER model, in which the discrete activity of network nodes is described by susceptible, excited and refractory states, representing a stylized neuron or neural population. In this case, the plasticity acts in a Hebbian-like fashion based on the functional connectivity (FC) derived from coactivation patterns of network nodes. The results confirm a correspondence between the two plasticity modalities, which speaks in favor of the dynamical implementation representing a biologically plausible mechanism through which topological reinforcement may take place in real systems, thus representing a fundamental model of the emergence of modular brain networks.

## Results

Starting from initial random configurations, we evolved networks according to the TR rule. Topological reinforcement was based on the TO between nodes of a network. At each rewiring step, a randomly selected node was connected to a non-neighbor with the highest TO, while pruning another link with random uniform probability, in order to preserve network density.

### Random networks evolve towards modular, small-world organization

TR reliably evolved synthetic random networks toward high modularity (Fig 1). Moreover, due to increased clustering, the final networks had a small-world organization (S1 Fig). The results were robust across multiple runs and multiple initial network realizations (S2 Fig). We also explored the effect of network size and density on the TR rule (Fig 1). The results were consistent, showing similar scaling curves across conditions, which speaks for the robustness of TR in generating modular networks.

**Fig 1.**
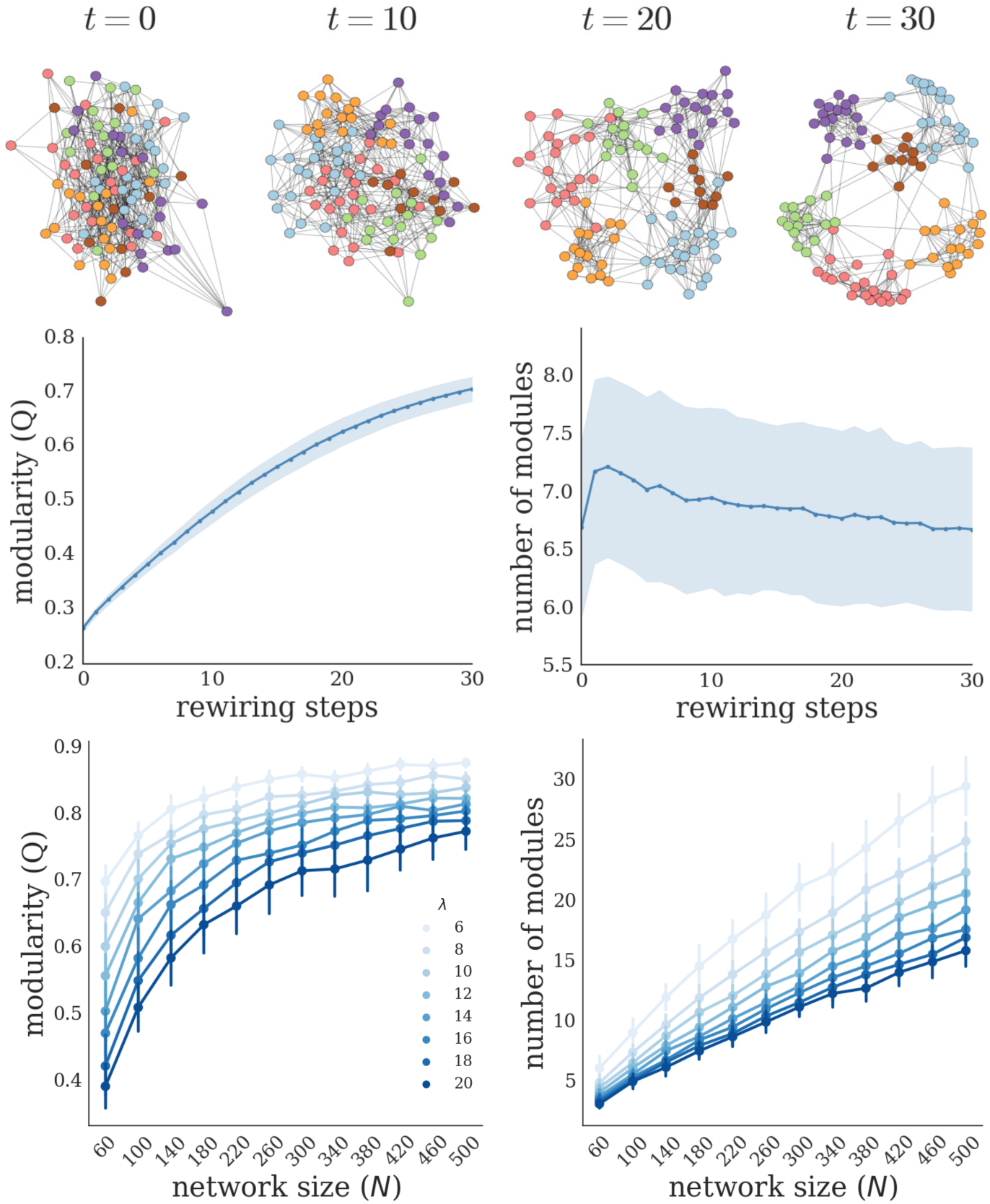
Emergence of modular network organization from topological reinforcement. (Top) Example of network evolution resulting from topological reinforcement, starting from a random network. Layouts are generated according to the Fruchterman-Reingold force-directed algorithm. Nodes are consistently colored according to the final modular structure. (Middle) Evolution of the modularity (Q); number of modules as a function of the number of rewiring steps (mean and standard deviation across 500 simulation runs). (Bottom) Final modularity (left) and number of modules detected (right) for different network sizes (*N*) and densities (*λ*) (mean and standard deviation across 50 independent graph realizations).

### Final network structure reflects initial network organization

TR appeared to amplify weak ‘proto-modules’ already present in the initial random graph. The similarities between the initial and final network structures were investigated in terms of Pearson correlation and partitions overlap between networks; see Methods section and Fig 2 for details.

**Fig 2.**
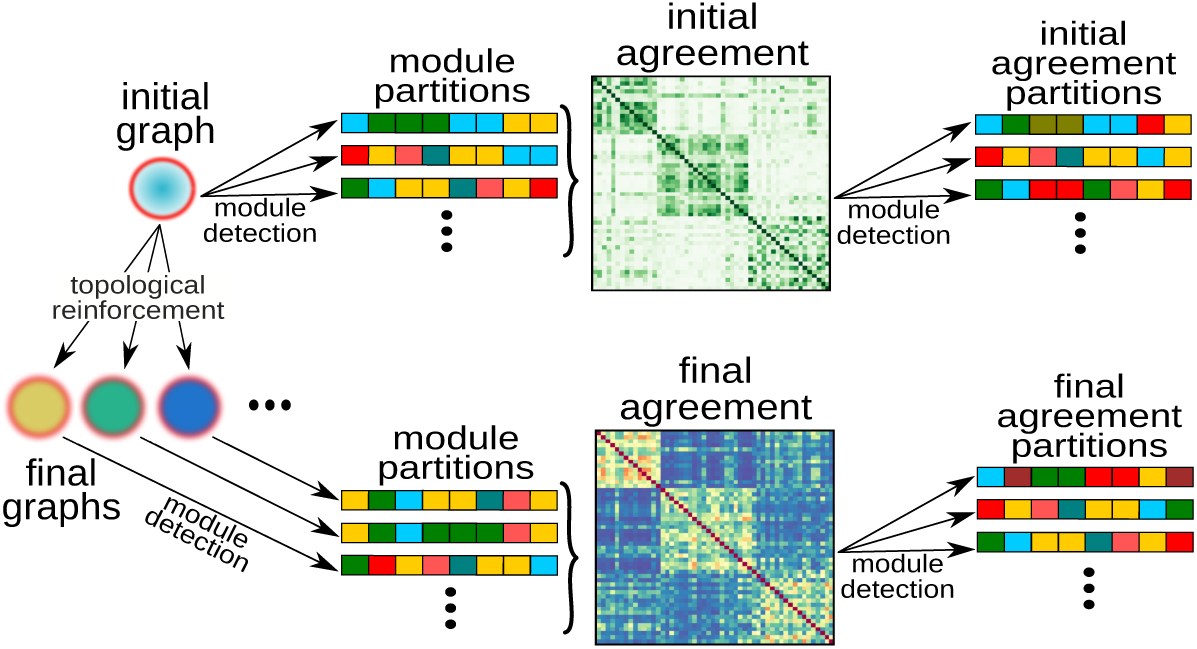
Modules agreement and ‘proto-modules’. Schematic representation of the procedure for probing the existence of ‘proto-modules’ in the initial graph and the relationship between initial and final network structure (see Methods for details).

Statistical analysis across multiple runs showed a significant similarity and partition overlap between the final graphs and the initial one (Fig 3 A). Moreover, the results also showed a consistent pattern of final modular organization (Fig 3 B). The module agreement of final networks across multiple runs (*P*) displayed pairs of nodes with high probability (beyond chance) to end up in the same module. Fig 3 B shows the mean intra-module density of the initial random graph according to different partitions. The distribution of the mean intra-module density according to the modules detected in the agreement *P* coincides fairly well with the mean intra-module density of the partitions detected on the graph itself. In contrast, intra-module density from partitions coming from a null model is centered around 0.1, that is, the graph density (i.e., probing density of randomly chosen groups of nodes).

**Fig 3.**
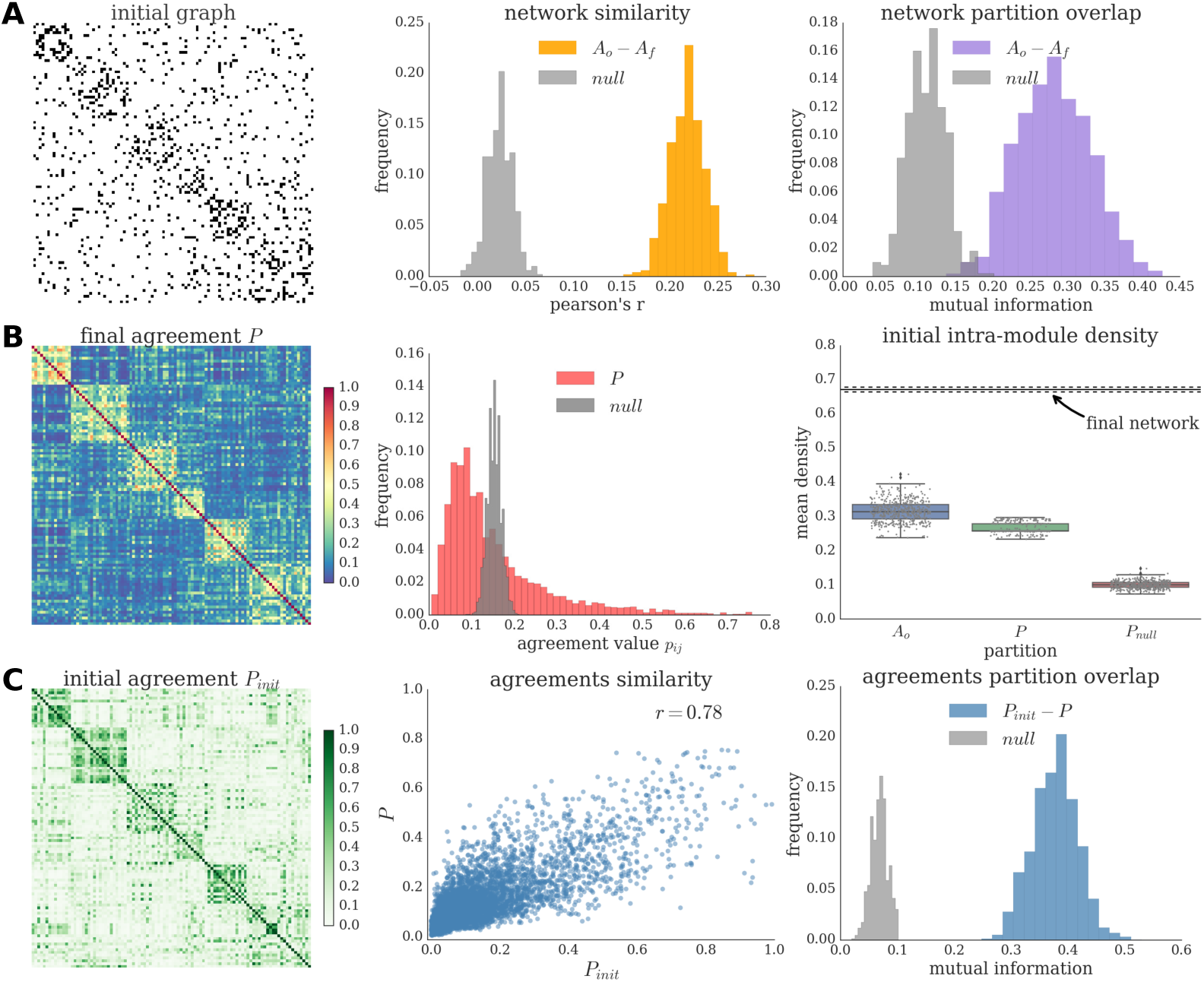
Relationship between initial and final network structures. (A) Initial adjacency matrix (left) reordered according to the modular partition of the agreement *P*. Similarity (middle) and partition overlap (right) between all pairs of initial and final networks, and the corresponding null distributions. (B) Agreement matrix across multiple runs (*P,* left) reordered according to its modular partition. Histogram of the *P* values and of the corresponding null model (middle). Distributions of the intra-modular density of the initial network (right). Average intra-module density of the initial network according to different types of module partitions. The procedure was repeated 500 times for each type of partition. As a reference, the mean intra-module density of the final network modules are also plotted (average and standard deviation). (C) Initial agreement matrix (*P*_*init*_, left) reordered according to the modular partition of *P*. Similarity (middle) and partition overlap (right) between *P*_*init*_ and *P* and the corresponding null distribution.

In the random graphs used as initial condition, no variations in link density are expected (since, by definition, connection probability is uniform for all pairs of nodes). Importantly, that is the case *on average across graph realizations*, but, due to stochastic variations and finite-size effect, individual graphs might contain groups of nodes with slightly higher density of edges than expected. We refer to these groups as ‘proto-modules’. In order to highlight these modules, a module detection algorithm was applied multiple times on the initial graph and a module agreement matrix was built (*P*_*init*_). The correspondence between the initial and final network structures is also evident comparing the final agreement *P* with its analogous on the initial graph *P*_*init*_ (Fig 3 C). The similarity (as measured by correlation) between both agreements is high. Additionally, we generated a set of partitions from *P* and another set of partitions from *P*_*init*_, and quantified the overlap between all possible pairs of partitions *P*_*init*_-*P*. We observed a significant overlap between the partitions from *P*_*init*_ and those from *P*. Furthermore, the results were robust across multiple initial network realizations (S3 Fig).

### Biological implementation of topological reinforcement

In the brain, the topological reinforcement may be implemented through various plausible activity-based models. We explored one such model, in which the activity of network nodes was described by discrete susceptible, excited and refractory states, the SER model, representing a stylized neuron or neural population. TR when transposed into biological context simply corresponds to the so-called Hebbian rule, where we substituted FC for TO, see Methods section for details.

In order to explore the FC-based rule and its relation to TR, we exploited an interesting feature of the SER model: for a given graph topology, the relationship between TO and FC varies according to the parameters of the model (transition probabilities *f, p* in the stochastic case and the initial conditions *e, s, r* in the deterministic case; for details, refer to [6]). Thus, after exhaustive evaluation of the possible constellations for each case, we found: first, that the FC-based rule was also able to generate a modular network structure. Importantly, a sufficiently high similarity (as measured by correlation) between TO and FC within the initial configuration was a necessary condition for modularity emergence, as illustrated by the sharp transition from the non-modular to the modular regime (Fig 4); second, the results produced by the FC-based plasticity were consistent with the ones from TR, both in terms of final network configurations and their module partitions (Fig 5). Fundamentally, this indicates that, provided the correlation between TO and FC is high enough, the Hebbian rule acts indirectly as topological reinforcement.

**Fig 4.**
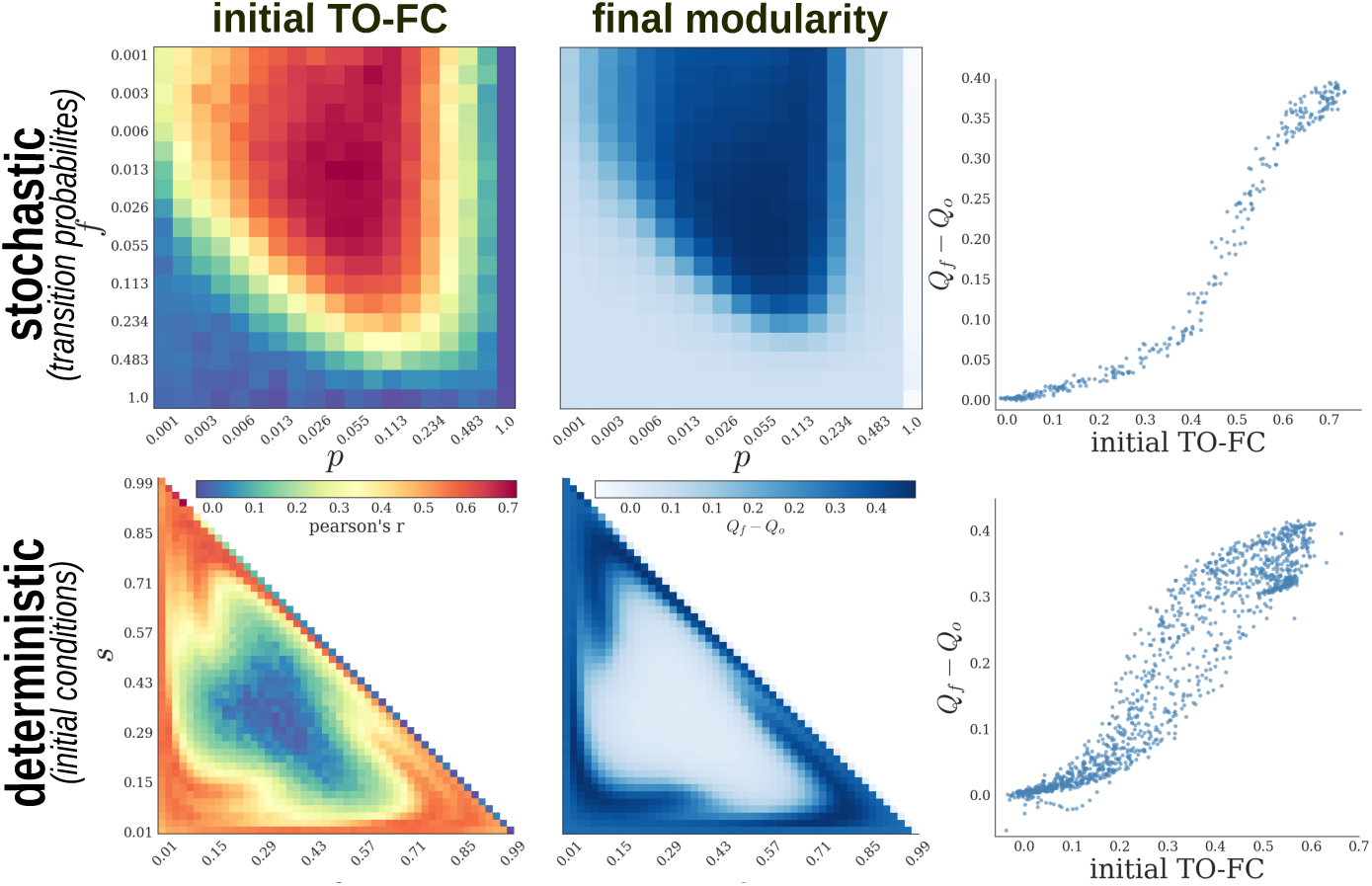
Biological implementation of the topological reinforcement. Parameter space exploration of the stochastic (top) and deterministic (bottom) SER model. Similarity (measured by correlation) between TO and FC in the initial graph (left), final modularity (middle) expressed as the difference between the mean final modularity value and the modularity of the initial random graph (across multiple (500) community detection). (Right) Scatter plot of the relationship between both quantities. Note logarithmic scale for the stochastic case.

**Fig 5.**
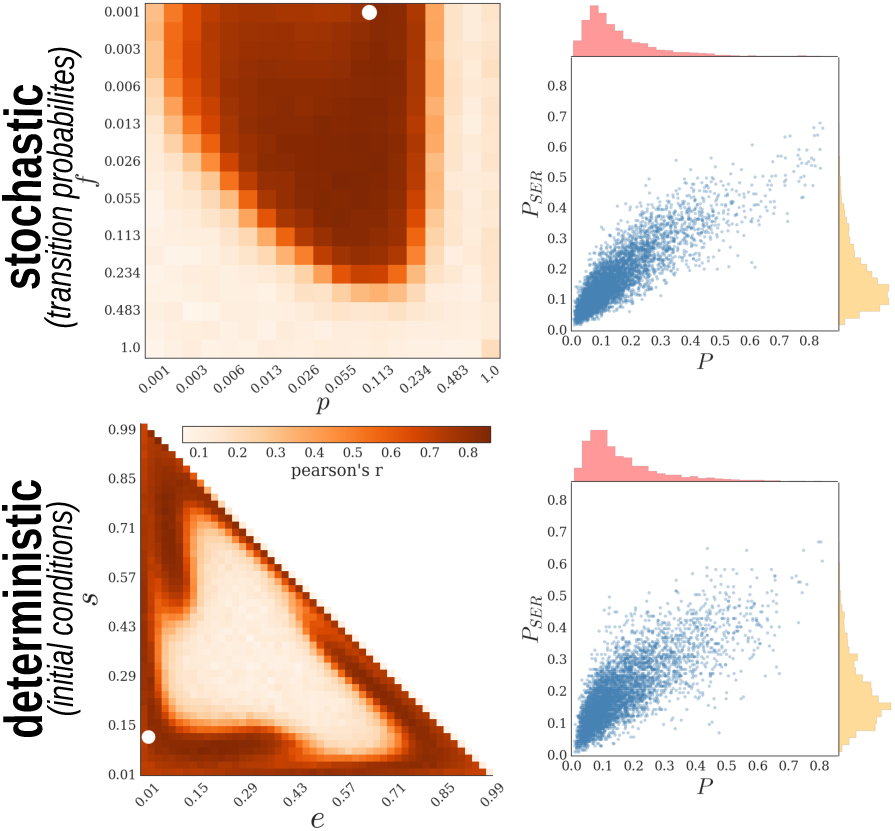
Correspondence between the topological reinforcement and the Hebbian rule. Similarity between *P* from the topological reinforcement and from the Hebbian rule using the stochastic (top) and deterministic (bottom) SER models. Pearson’s correlation coefficient was computed to summarize the similarity between both rules across the parameter spaces. Scatter plots represent the relationship for a selected setting (white dots in the heatmaps).

## Discussion

The importance of segregation in the brain is supported by numerous studies [1,34]. However, there is a lack of general mechanisms explaining the emergence of brain modularity. In the present study, we proposed an explicit mechanism of reshaping local neighbourhoods through topological reinforcement that might act as a fundamental principle underlying the emergence of modules in brain networks.

Given accumulated evidence that global network properties can systematically affect the composition of local network structure such as motifs [31-33], we propose a complementary bottom-up approach that is acting locally in order to shape global features. Our proposed mechanism is in line with empirical data where ‘homophily’ appears as an essential feature of brain connectivity. At the micro scale, it has been shown that the probability to find a connection between a pair of neurons is proportional to the number of their shared neighbours [35]; while, at the macro scale, the strength of connections between brain regions tends to be the higher the more similar their connectivity profiles are [36].

Our results show that local reinforcement reliably and robustly produces modular network architectures over time, accompanied by the small-world property. Additionally, the final modular organization of the networks seems to correspond to groups of nodes in the initial networks that have higher than average connection density. As such, our rewiring mechanism acts as an amplification of these proto-modules, similarly to a previously reported effect in weak modular weighted networks evolving under a Hebbian rule based on chaotic maps synchronization [37].

We extended the framework of topological reinforcement by introducing a plausible biological implementation. Our dynamical model choice, the SER model, offers the advantage of capturing essential characteristics of stylized neuronal activity while being easily tractable. This minimalistic excitable network model has a rich history across disciplines and in particular in neurosciences [38-42], where it can capture non-trivial statistical features of brain activity patterns [43,44]. This model has also been used to study the impact of network topology, such as modules, hubs and cycles, on network activity patterns [5,44,45]. A relative-threshold variant (requiring a certain percentage of a node’s neighbors to be active, in order to activate the node) was explored in [46] and [47]. The deterministic limit of the model (*p* →1, *f* →0) has been analyzed in [48] and in much detail in [6].

In the biological implementation, the topological reinforcement rule was reformulated by using functional connectivity as a surrogate of TO. The results were consistent with TR, indicating that the biological implementation appears to indirecly act at the topological level. In other words, the FC served as a proxy of TO, and therefore Hebbian reinforcement led indirectly and ultimately to the topological reinforcement of a modular network organizatiion. The explanation for this finding is based on the fact that, for suitable dynamical regimes and structural architectures, FC is positively correlated with TO in excitable networks [6], which is intuitive if one considers that common inputs may promote correlations. Our results are in line with recent theoretical work on the contribution of specific network motifs to higher order network organization, in which the reinforcement of connections between neurons receiving common inputs led to the formation of self-connected assemblies [49]. Hence, our Hebbian plasticity scenario exploited the correspondence between TO and FC as it could be observed with the exploration of different SER parameter constellations. These parameters promote different relations between TO and FC, and we found that such a dependence systematically predicted the emergence (or not) of modular networks.

Previous computational studies have shown that evolutionary algorithms of network connectivity optimizing, for example, functional complexity (defined as balance between segregation and integration) can lead to modular network formation [50]. Such findings point to the relevance of modularity as a crucial organization principle underlying complex functional brain processes. Nevertheless, these models do not provide a biologically interpretable and implementable mechanism, since the explicit global optimization function (functional complexity) which might be relevant at the evolutionary scale, cannot be directly interpreted as biological mechanisms shaping brain connectivity.

In the sense of biological plausibility, activity-based plasticity models (e.g., based on Hebbian plastiticy) constitute a more directly interpretable approach. Previous studies have used a variety of neural activity models ranging from abstract representations, such as chaotic maps [51] and phase oscillators [52], to more physiologically realistic models, such as neural masses [53] and spiking neuron [54] models. In general, Hebbian reinforcement led to the formation of modular architectures, consistent with our results for the excitable model. The open question for this type of models concerns the specific underlying topological changes that they promote, since these studies focus on the implementation of the phenomenon (based on the activity) and not on the algorithmic level (the topological dimension). Both explanatory levels interact in non-trivial ways. Indeed some of these models even showed that final topological features (e.g., number of modules) might purely depend on properties of the dynamical model [37]. In other words, they do not provide insights about a general mechanism specifying which topological changes might be necessary for the emergence of modular structure.

An alternative approach is provided by generative models, where typically an objective function governs the insertion of links and/or nodes during simulations. Recent works has shown that including homophily as a factor to determine connection probability (and after proper data-driven parameter tuning) makes it possible to account for a great deal of functional [55] as well as structural [17] topological features of real large-scale brain networks. While these studies provide a valuable basis for confirming the importance of TO as an essential feature and reducing the dimensionality of brain connectivity by few model parameters, disentangling the mechanistic nature of the phenomena (e.g., modularity emergence) turns out to be non-trivial, since information about the final state is explicitly built-in in the generative model already. Moreover, how the generative function is actually implemented in real systems is out of the scope of this kind of modeling.

In summary, as expected for any modeling approach, there exists a complementarity between generative and activity-based models, in which they trade-off description and mechanistic explanations and a gap remains for explaining on how they link to each other. Our contribution represents an attempt to address this gap; first, by providing an explicit topological mechanism of module formation (generative mechanism); second, by trying to reconcile such an abstract level of analysis with the biological implementation, by means of an activity-based variation of the model.

The present results are subject to several methodological considerations, for example, the absence of geometric factors in the model. Although the brain is a spatially embedded system and physical constrains play a fundamental role constraining brain connectivity [11], the focus of our study was on the topological determinants of the formation of brain networks. We aimed at avoiding the situation in which geometric constrains, such as the distance-dependent probability of connection used in previous studies [56], introduce already by themselves a clustered connectivity, thus potentially overriding the changes based on the topology itself. Regarding the plausible biological implementation, we chose a simple abstract model for computational tractability. It would be interesting to compare our framework with more biologically realistic dynamical models, such as networks of spiking neurons.

## Conclusions

Our findings suggest a selective reinforcement of the topological overlap as a plausible mechanism by which brain networks evolve toward a modular organization. Moreover, under appropriate conditions, functional connectivity might act as a proxy, or a dynamical representation, of TO. Thus, biological-inspired plasticity rules, such as the Hebbian rule, indirectly promote modularity. To our knowledge, these findings constitute a first topologically mechanistic explanation of modules formation in complex brain networks and its link to a physiologically plausible realization. Despite of the simplicity of our framework, we trust it to carry a conceptual value that contributes to the long challenging path of understanding the fundamental principles of brain organization.

## Methods

### Networks

We considered synthetic undirected networks without self-connections of size *N* = 100 nodes and average connectivity *λ* = 10 (equivalently, a density of 0.1). The networks were represented by a symmetric adjacency matrix A, where *a*_*ij*_ = 1, if nodes *i* and *j* are connected, 0 otherwise. Initial networks were generated according to the classical Erdös-Rényi model [57].

We explored the robustness of the plasticity rule across various network realizations and multiple runs (using the same initial network). We generated 100 synthetic random initial graphs and performed 500 runs for each of them. In order to study the scaling properties of our model, we also evaluated graphs with different average connectivity (between 6 and 20 by step of 2) and size (between 60 and 500 by step of 40).

### Topological reinforcement

Topological reinforcement was based on the topological overlap measure. TO represents the neighborhoods’ similarity of a pair of nodes by counting their number of common neighbors [23]:

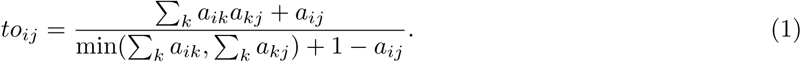

At each rewiring step, the rule connects a randomly selected node that is neither disconnected nor fully connected with a non-neighbor with the highest TO, while pruning another link with uniform probability, hence preserving graph density.

For computational efficiency, the rewiring was applied by inserting simultaneously one link on 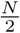 random different nodes at each step, and pruning the same number of links at random, so that 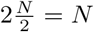 links were reallocated at each rewiring step, with statistically equivalent results as when only two links (one insertion, one pruning) per step were modified.

In order to compare the results across different graph sizes and densities, we computed the length of each run, *r*, by fixing the average number of rewiring per link, 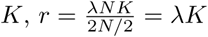. Throughout the manuscript *K*= 3.

### Excitable model

We used a three-state cellular automaton model of excitable dynamics, the SER model. The activity evolves according to the following synchronous transition rules:

- S → E, if at least one neighbour is excited; or with probability *f (*spontaneous activation);
- E → R;
- R → S, with probability *p (*recovery).

In the deterministic SER scenario, i.e., *f* = 0 and *p* = 1, for each network and initial condition setting, the activity time windows consisted of 5 000 runs of 30 time steps each and FC was averaged over runs. The initial conditions were randomly generated, covering the full space of possible proportions of states.

In the stochastic SER scenario, i.e., *f >* 0 and *p <* 1, for each parameter setting (*f, p*), the activity time window consisted of one run of 50 000 time steps. The initial conditions were randomly generated with a proportion of 0.1 nodes excited, while the remaining nodes were equipartitioned into susceptible and refractory states.

### Functional connectivity

To analyze the pattern of excitations in the SER model, we computed the number of joint excitations for all possible pairs of nodes. The outcome matrix is the so-called coactivation matrix, a representation of the functional connectivity of the nodes:

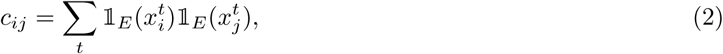

where 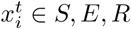 being the state of node *i* at time *t*, and 1_*E*_ the indicator function of state E. FC was then normalized to scale values between 0 and 1:

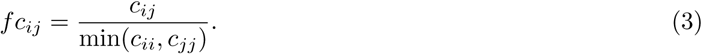

### Biological implementation-Hebbian rule

When transposing the topological reinforcement into a biological context, using a plausible model of brain dynamics, it turns out that the rule corresponds to the well known Hebbian rule, where we substituted FC for TO.

We used SER model for a simulation run after which FC was derived and the rewiring was applied: a random node was selected and connected to a non-neighbor node with maximum FC, while a link was selected randomly with uniform probability and pruned. As for the topological reinforcement and for computational efficiency, the rewiring was applied simultaneously on 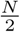 different nodes at each step. In order to keep the final networks comparable, the total number of rewiring steps was the same for both plasticity modalities, as defined above.

According to the SER scenario, stochastic or deterministic, we evaluated the model for different parameter constellations or initial conditions, respectively. For one initial graph, we studied each possible combination of parameter constellation/initial condition by performing 150 simulation runs and the final graph measures were averaged across runs.

## Network analysis

Synthetic graph realizations, basic graph properties (clustering, path length, small-world), community detection, matrix reordering and graph layouts were performed using the Brain Connectivity Toolbox [58] (Python version 0.5.0; github.com/aestrivex/bctpy) and NetworkX [59]. For a given graph, communities were extracted by means of the Louvain algorithm that attempts to maximize the modularity of the network, using the so-called Q value [60].

Similarity between networks and agreements was assessed by means of the Pearson correlation between their connectivity matrices. Overlap between partitions was probed based on the normalized mutual information between the communities [61].

### Module agreement and ‘proto-modules’

From a given initial network, multiple simulation runs (500) were performed and the community detection algorithm was applied on each final graph to find a partition of the nodes into communities. Then, an agreement matrix *P* was computed across all final partitions, where *p*_*ij*_ quantifies the frequency with which nodes *i* and *j* belonged to the same community across partitions. Finally, the community detection algorithm was applied 100 times on *P,* yielding a representative set of final partitions of the nodes into non-overlapping communities given an initial graph (Fig 2).

In order to probe the structure of each initial graph and find potential ‘proto-modules’, we applied the community detection on the initial graph. Due to the weak signal of random graphs, the stochasticity and associated degeneracy of classical community detection algorithms, a consensus clustering was employed to generate stable solutions. For each random initial graph, the community detection algorithm was applied 500 times, then a agreement matrix was computed, named *P*_*init*_, and finally the community detection algorithm was applied 100 times on this agreement matrix yielding a representative set of (stable) partitions of the initial graph (Fig 2).

### Statistical assessments

In order to assess the significance of the results, null network models were generated. When comparing networks in terms of similarity (by Pearson correlation),a null model was generated by randomly rewiring a given graph (once per link), while preserving the degree distribution [62]. Two null models where used when comparing networks in terms of partition overlap. For comparison of individual runs (initial vs. final structures or initial vs. final agreements), we simply used a rewired initial graph as explained above instead of the actual one that was used as initial condition for the run. As null model for the comparison of agreement matrices, a null agreement *P*_*null*_ was constructed by first shuffling the individual partitions (i.e., conserving the number of modules and their sizes, but randomly altering the nodes affiliation) and then computing the agreement across them. Thus, such a null model generates the expected distribution of agreement values that would occur purely by chance for a given number of nodes and modules of given sizes.

## Supporting information

**S1 Fig.**
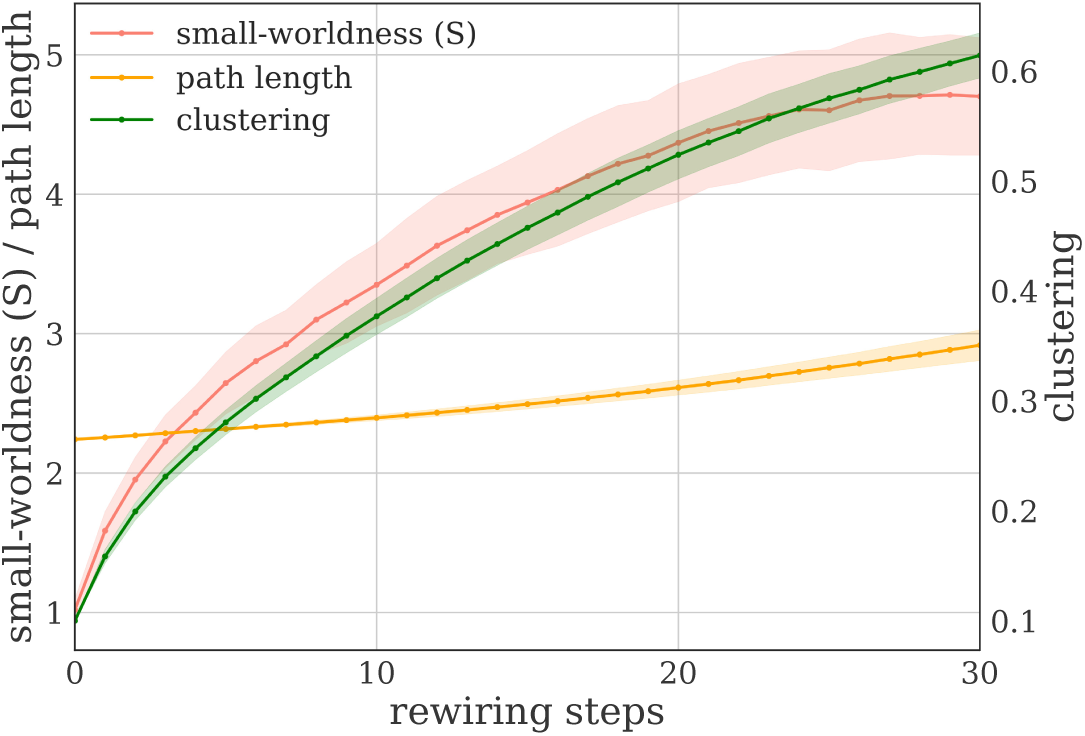
Evolution of the mean clustering coefficient, characteristic path length and small-world index (S). Results are expressed as mean and standard deviation across 500 simulation runs as a function of the number of rewiring steps.

**S2 Fig.**
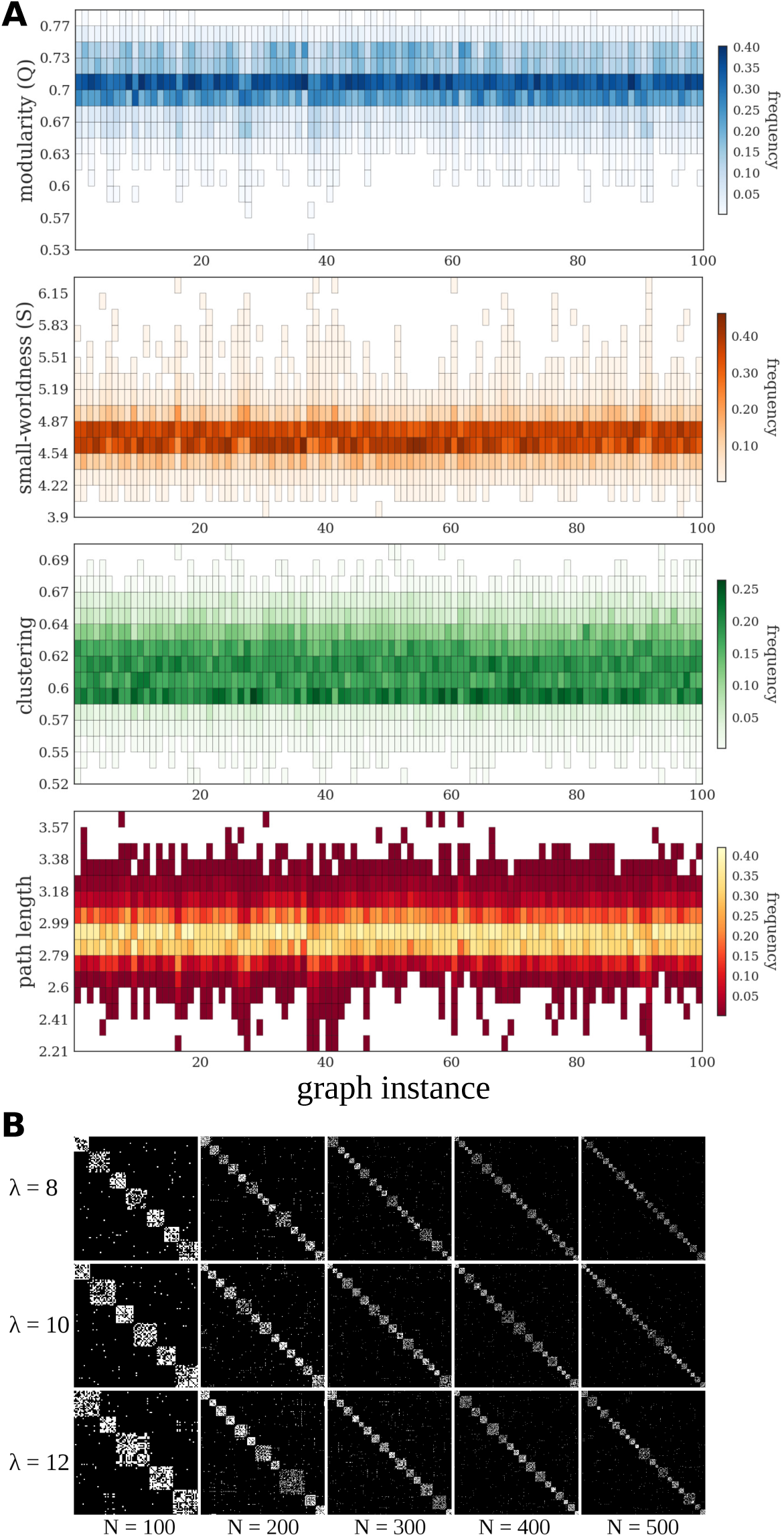
Robustness of results for different initial random graph instances, final networks characteristics. (A) The heat maps show a summary of the results for 100 different initial random graph instances used as initial condition. For each one, 500 simulation runs were performed. For each final graph, modularity (Q), characteristic path length, mean clustering coefficient and and small-worldness coefficient (S) were computed. Each column of the heat map represents the distribution of values obtained across 500 simulations carried out with the same initial random graph instance. (B) Examples of final adjacency matrices of individual runs with different network sizes (*N*) and densities (*λ*).

**S3 Fig.**
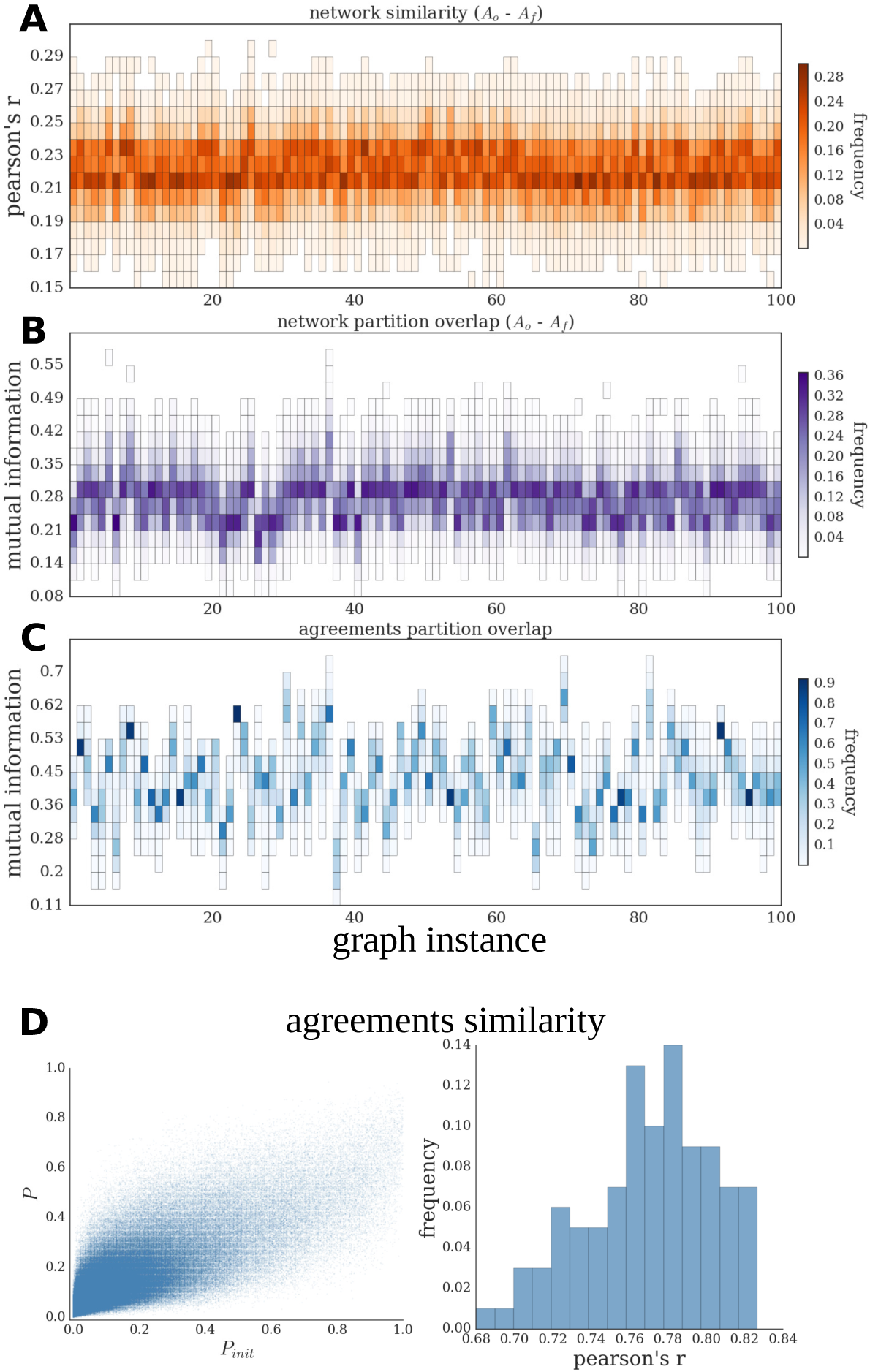
Robustness of results for different initial random graph instances, correspondence between initial and final networks. (A-C) The heat maps show a summary of the results for 100 different initial random graph instances used as initial condition. Each column of the heat maps represents the distribution of values obtained across 500 simulations carried out with the same initial random graph instance.(A) Distribution of the correlation values between all pairs of initial and final networks. (B) Partition overlap between all pairs of initial and final networks (quantified as normalized mutual information, NMI). (C) Partition distance between initial (*P*_*init*_) and final (*P*) agreements. (D) Similarity between *P*_*init*_ and *P* agreements, the scatter plot (left) shows the values over 100 different random graph instances used as initial condition, and the histogram (right) represents the Pearson’s correlation coefficient between all pairs of *P*_*init*_-*P*.

## Acknowledgements

FD was supported by the Deutscher Akademischer Austauschdienst (DAAD) as well as the Deutsche Forschungsgemeinschaft (DFG) grant SFB 936/A1. CCH was supported by DFG grants HI 1286/5-1, HI 1286/6-1, HI 1286/7-1, SFB 936/A1, Z3 and TRR 169/A2. MTH acknowledges support from DFG grant HU 937/7. AM acknowledges support from the DFG grant SFB 936/Z3. The funders had no role in study design, data collection and analysis, decision to publish, or preparation of the manuscript.

## References

1. Sporns O, Betzel RF. Modular Brain Networks. Annual review of psychology. 2016;67(1):613–40. doi:10.1146/annurev-psych-122414-033634.

2. Rosvall M, Bergstrom CT. Maps of random walks on complex networks reveal community structure. Proceedings of the National Academy of Sciences. 2008;105(4):1118–1123. doi:10.1073/pnas.0706851105.

3. Arenas A, Díaz-Guilera A, Péerez-Vicente CJ. Synchronization Reveals Topological Scales in Complex Networks. Phys Rev Lett. 2006;96:114102. doi:10.1103/PhysRevLett.96.114102.

4. Zhou C, Zemanováa L, Zamora G, Hilgetag CC, Kurths J. Hierarchical Organization Unveiled by Functional Connectivity in Complex Brain Networks. Phys Rev Lett. 2006;97:238103. doi:10.1103/PhysRevLett.97.238103.

5. Müller-Linow M, Hilgetag CC, Hütt MT. Organization of Excitable Dynamics in Hierarchical Biological Networks. PLOS Computational Biology. 2008;4(9):1–15. doi:10.1371/journal.pcbi.1000190.

6. Messé A, Hütt MT, Hilgetag CC. Toward a theory of coactivation patterns in excitable neural networks. PLOS Computational Biology. 2018;14(4):1–19. doi:10.1371/journal.pcbi.1006084.

7. Hilgetag C, Burns GAPC, O’Neill MA, Scannell JW, Young MP. Anatomical connectivity defines the organization of clusters of cortical areas in the macaque and the cat. Philosophical Transactions of the Royal Society of London B: Biological Sciences. 2000;355(1393):91–110. doi:10.1098/rstb.2000.0551.

8. Kaiser M, Hilgetag C. Optimal hierarchical modular topologies for producing limited sustained activation of neural networks. Frontiers in Neuroinformatics. 2010;4:8. doi:10.3389/fninf.2010.00008.

9. Wang SJ, Zhou C. Hierarchical modular structure enhances the robustness of self-organized criticality in neural networks. New Journal of Physics. 2012;14(2):023005.

10. Baum GL, Ciric R, Roalf DR, Betzel RF, Moore TM, Shinohara RT, et al. Modular Segregation of Structural Brain Networks Supports the Development of Executive Function in Youth. Current Biology. 2017;27(11):1561–1572.e8. doi:https://doi.org/10.1016/j.cub.2017.04.051.

11. Henderson JA, Robinson Pa. Using Geometry to Uncover Relationships Between Isotropy, Homogeneity, and Modularity in Cortical Connectivity. Brain Connectivity. 2013;3(4):423–437. doi:10.1089/brain.2013.0151.

12. Gómez-Robles A, Hopkins WD, Sherwood CC. Modular structure facilitates mosaic evolution of the brain in chimpanzees and humans. Nature communications. 2014;5:4469. doi:10.1038/ncomms5469.

13. Ellefsen KO, Mouret JB, Clune J. Neural Modularity Helps Organisms Evolve to Learn New Skills without Forgetting Old Skills. PLoS Computational Biology. 2015;11(4):1–24. doi:10.1371/journal.pcbi.1004128.

14. Betzel RF, Bassett DS. Generative models for network neuroscience: prospects and promise. Journal of The Royal Society Interface. 2017;14(136). doi:10.1098/rsif.2017.0623.

15. Kaiser M, Hilgetag CC. Development of multi-cluster cortical networks by time windows for spatial growth. Neurocomputing. 2007;70(10):1829–1832. doi:https://doi.org/10.1016/j.neucom.2006.10.060.

16. Bauer R, Kaiser M. Nonlinear growth: an origin of hub organization in complex networks. Open Science. 2017;4(3). doi:10.1098/rsos.160691.

17. Betzel RF, Avena-Koenigsberger A, Goñi J, He Y, de Reus MA, Griffa A, et al. Generative models of the human connectome. NeuroImage. 2016;124:1054–1064. doi:https://doi.org/10.1016/j.neuroimage.2015.09.041.

18. Gong P, van Leeuwen C. Emergence of scale-free network with chaotic units. Physica A: Statistical Mechanics and its Applications. 2003;321(3):679 – 688. doi:https://doi.org/10.1016/S0378-4371(02)01735-1.

19. Abbott LF, Nelson SB. Synaptic plasticity: taming the beast. Nature Neuroscience. 2000;3:1178 EP –.

20. Effenberger F, Jost J, Levina A. Self-organization in Balanced State Networks by STDP and Homeostatic Plasticity. PLOS Computational Biology. 2015;11(9):1–30. doi:10.1371/journal.pcbi.1004420.

21. Stone D, Tesche C. Topological dynamics in spike-timing dependent plastic model neural networks. Frontiers in Neural Circuits. 2013;7:70. doi:10.3389/fncir.2013.00070.

22. Jarman N, Steur E, Trengove C, Tyukin IY, van Leeuwen C. Self-organisation of small-world networks by adaptive rewiring in response to graph diffusion. Scientific Reports. 2017;7(1):13158. doi:10.1038/s41598-017-12589-9.

23. Ravasz E, Somera AL, Mongru DA, Oltvai ZN, Barabási AL. Hierarchical Organization of Modularity in Metabolic Networks. Science. 2002;297(5586):1551–1555. doi:10.1126/science.1073374.

24. Bass JIF, Diallo A, Nelson J, Soto JM, Myers CL, Walhout AJ. Using networks to measure similarity between genes: association index selection. Nature methods. 2013;10(12):1169.

25. Hilgetag CC. Mathematical approaches to the analysis of neural connectivity in the mammalian brain. University of Newcastle upon Tyne; 1999.

26. Hilgetag CC, Kötter R, Stephan KE, Sporns O. Computational methods for the analysis of brain connectivity. In: Computational neuroanatomy. Springer; 2002. p. 295–335.

27. Sporns O. Graph theory methods for the analysis of neural connectivity patterns. In: Neuroscience databases. Springer; 2003. p. 171–185.

28. Li A, Horvath S. Network neighborhood analysis with the multi-node topological overlap measure. Bioinformatics. 2006;23(2):222–231.

29. Milo R, Shen-Orr S, Itzkovitz S, Kashtan N, Chklovskii D, Alon U. Network motifs: simple building blocks of complex networks. Science. 2002;298(5594):824–827.

30. Milo R, Itzkovitz S, Kashtan N, Levitt R, Shen-Orr S, Ayzenshtat I, et al. Superfamilies of evolved and designed networks. Science. 2004;303(5663):1538–1542.

31. Vazquez A, Dobrin R, Sergi D, Eckmann JP, Oltvai Z, Barabási AL. The topological relationship between the largescale attributes and local interaction patterns of complex networks. Proceedings of the National Academy of Sciences. 2004;101(52):17940–17945.

32. Fretter C, Müller-Hannemann M, Hütt MT. Subgraph fluctuations in random graphs. Physical Review E. 2012;85(5):056119.

33. Reichardt J, Alamino R, Saad D. The interplay between microscopic and mesoscopic structures in complex networks. PloS one. 2011;6(8):e21282.

34. Wig GS. Segregated Systems of Human Brain Networks. Trends in Cognitive Sciences. 2017;21(12):981 – 996. doi:https://doi.org/10.1016/j.tics.2017.09.006.

35. Perin R, Berger TK, Markram H. A synaptic organizing principle for cortical neuronal groups. Proceedings of the National Academy of Sciences. 2011;108(13):5419–5424. doi:10.1073/pnas.1016051108.

36. Goulas A, Schaefer A, Margulies DS. The strength of weak connections in the macaque cortico-cortical network. Brain Structure and Function. 2015;220(5):2939–2951. doi:10.1007/s00429-014-0836-3.

37. Yuan WJ, Zhou C. Interplay between structure and dynamics in adaptive complex networks: Emergence and amplification of modularity by adaptive dynamics. Phys Rev E. 2011;84:016116. doi:10.1103/PhysRevE.84.016116.

38. Bak P, Chen K, Tang C. A forest-fire model and some thoughts on turbulence. Physics Letters A. 1990;147(5):297 – 300. doi:https://doi.org/10.1016/0375-9601(90)90451-S.

39. Anderson R, May RM. Infectious Diseases of Humans: Dynamics and Control. Oxford: Oxford University Press; 1992.

40. Drossel B, Schwabl F. Self-organized critical forest-fire model. Phys Rev Lett. 1992;69:1629–1632. doi:10.1103/PhysRevLett.69.1629.

41. Kinouchi O, Copelli M. Optimal dynamical range of excitable networks at criticality. Nature Physics. 2006;2:348 EP –.

42. Furtado LS, Copelli M. Response of electrically coupled spiking neurons: A cellular automaton approach. Phys Rev E. 2006;73:011907. doi:10.1103/PhysRevE.73.011907.

43. Haimovici A, Tagliazucchi E, Balenzuela P, Chialvo DR. Brain Organization into Resting State Networks Emerges at Criticality on a Model of the Human Connectome. Phys Rev Lett. 2013;110:178101. doi:10.1103/PhysRevLett.110.178101.

44. Messé A, Hütt MT, König P, Hilgetag CC. A closer look at the apparent correlation of structural and functional connectivity in excitable neural networks. Scientific reports. 2015;5:7870. doi:10.1038/srep07870.

45. Garcia GC, Lesne A, Hilgetag CC, Hütt MT. Role of long cycles in excitable dynamics on graphs. Phys Rev E. 2014;90:052805. doi:10.1103/PhysRevE.90.052805.

46. Hütt MT, Jain MK, Hilgetag CC, Lesne A. Stochastic resonance in discrete excitable dynamics on graphs. Chaos, Solitons & Fractals. 2012;45(5):611 – 618. doi:https://doi.org/10.1016/j.chaos.2011.12.011.

47. Fretter C, Lesne A, Hilgetag CC, Hütt MT. Topological determinants of self-sustained activity in a simple model of excitable dynamics on graphs. Scientific Reports. 2017;7:42340 EP –.

48. Garcia GC, Lesne A, Hütt Mt, Hilgetag CC. Building Blocks of Self-Sustained Activity in a Simple Deterministic Model of Excitable Neural Networks. Frontiers in Computational Neuroscience. 2012;6(August):50. doi:10.3389/fncom.2012.00050.

49. Ravid Tannenbaum N, Burak Y. Shaping Neural Circuits by High Order Synaptic Interactions. PLOS Computational Biology. 2016;12(8):1–27. doi:10.1371/journal.pcbi.1005056.

50. Sporns O, Tononi G, Edelman GM. Connectivity and complexity: the relationship between neuroanatomy and brain dynamics. Neural Networks. 2000;13(8):909–922. doi:https://doi.org/10.1016/S0893-6080(00)00053-8.

51. van den Berg D, van Leeuwen C. Adaptive rewiring in chaotic networks renders small-world connectivity with consistent clusters. EPL (Europhysics Letters). 2004;65(4):459.

52. Gleiser PM, Zanette DH. Synchronization and structure in an adaptive oscillator network. The European Physical Journal B - Condensed Matter and Complex Systems. 2006;53(2):233–238. doi:10.1140/epjb/e2006-00362-y.

53. Stam CJ, Hillebrand A, Wang H, Van Mieghem P. Emergence of Modular Structure in a Large-Scale Brain Network with Interactions between Dynamics and Connectivity. Frontiers in Computational Neuroscience. 2010;4:133. doi:10.3389/fncom.2010.00133.

54. Kwok HF, Jurica P, Raffone A, van Leeuwen C. Robust emergence of small-world structure in networks of spiking neurons. Cognitive Neurodynamics. 2006;1(1):39. doi:10.1007/s11571-006-9006-5.

55. Vértes PE, Alexander-Bloch AF, Gogtay N, Giedd JN, Rapoport JL, Bullmore ET. Simple models of human brain functional networks. Proceedings of the National Academy of Sciences. 2012;109(15):5868–5873. doi:10.1073/pnas.1111738109.

56. Jarman N, Trengove C, Steur E, Tyukin I, van Leeuwen C. Spatially constrained adaptive rewiring in cortical networks creates spatially modular small world architectures. Cognitive Neurodynamics. 2014;8(6):479–497. doi:10.1007/s11571-014-9288-y.

57. Erdős P, Rényi A. On Random Graphs I. Publicationes Mathematicae (Debrecen). 1959;6:290–297.

58. Rubinov M, Sporns O. Complex network measures of brain connectivity: Uses and interpretations. NeuroImage. 2010;52(3):1059 – 1069. doi:https://doi.org/10.1016/j.neuroimage.2009.10.003.

59. Hagberg AA, Schult DA, Swart PJ. Exploring Network Structure, Dynamics, and Function using NetworkX. In: Varoquaux G, Vaught T, Millman J, editors. Proceedings of the 7th Python in Science Conference. Pasadena, CA USA; 2008. p. 11 – 15.

60. Blondel VD, Guillaume JL, Lambiotte R, Lefebvre E. Fast unfolding of communities in large networks. Journal of Statistical Mechanics: Theory and Experiment. 2008;2008(10):P10008.

61. Meilă M. Comparing clusterings—an information based distance. Journal of Multivariate Analysis. 2007;98(5):873 – 895. doi:https://doi.org/10.1016/j.jmva.2006.11.013.

62. Maslov S, Sneppen K. Specificity and Stability in Topology of Protein Networks. Science. 2002;296(5569):910–913. doi:10.1126/science.1065103.

